# Microprobe-XRF assessment of nutrient distribution in soybean, cowpea, and kidney bean seeds: a Fabaceae family case study

**DOI:** 10.1101/2022.09.23.509231

**Authors:** Gabriel Sgarbiero Montanha, Sara Luiza Zachi Romeu, João Paulo Rodrigues Marques, Lívia Araújo Rohr, Eduardo de Almeida, André Rodrigues dos Reis, Francisco Scaglia Linhares, Sabrina Sabatini, Hudson Wallace Pereira de Carvalho

**Author notes:** Correspondence: Associate Prof. Hudson W. P. Carvalho.

## Abstract

The present study explored the microprobe X-ray fluorescence spectroscopy (µ-XRF) for quantitative and space-resolved distribution of macro, *i*.*e*., K, P, S, and Ca, and micronutrients, *i*.*e*., Fe, Zn, and Mn elemental composition in the cross-sectioned seeds of cowpea (*Vigna unguiculata* L.), kidney bean (*Phaseolus vulgaris* L.), and soybean *(Glycine max* L.) seeds, important agricultural species within the Fabaceae family. It unveils that both macro and micronutrients were heterogeneously distributed across seed tissues. Most of the P and S, Fe, Zn, and Mn were mainly found at the embryo axis tissues in all three Fabaceae species, whereas K was spread along the cotyledon and Ca was mostly observed trapped at the seed coat region. Furthermore, the Pearson correlation coefficient revealed a strong spatial correlation between P and S, and K and S in cowpea and soybean seed tissues, whereas Zn and Mn association was also recorded. Therefore, the µ-XRF technique demonstrates to be an important tool for assessing seed nutrient distribution, thus subsidizing the understanding of the physiological role of nutrients in seeds and fostering innovative approaches for nutrient supply and biofortification.

## 1. INTRODUCTION

The Fabaceae (Leguminosae) is the third-largest family of flowering plants [1], encompassing *ca*. 800 genera and 20,000 described species [2], some of which are important species for livestock feed and human food, such as peanuts (*Arachis hypogaea*), peas (*Pisum sativum*), common bean (*Phaseolus vulgaris*), chickpea (*Cicer arietinum*), cowpea (*Vigna unguiculata*), lentil (*Lens culinaris*), and soybean (*Glycine max (L*.*) Merrill*). Notably, the legume seeds are a remarkable source of carbohydrates, proteins, and oil [3,4] and only soybean seeds account for more than 1:2 of the seed-based oil and 2:3 of all the protein meals currently produced worldwide [5].

As far as the storage nutrients play crucial roles in seed germination and seedling establishment [6,7], the mineral composition of seeds has been widely investigated aiming at understanding either their effects on the development of plants under stressful conditions, *e*.*g*., drought or salinity stress or mechanisms to boost its nutritional quality [7–10].

On the other hand, space-resolved nutrient localization revealed that nutrient distribution varies across seed tissues. Particularly, P, S, and Ca were mainly found in the embryonic regions, whereas Ca was found in seed coat and hilum regions of *Crotalaria ochroleuca* and *Arabidopsis thaliana* seeds [11,12], whereas beneficial elements, such as Se and Ni, respectively, can be found in the embryo and cotyledon tissues of cowpea (*Vigna unguiculata* L.), buckwheat (*Fagopyrum esculentum*), and soybean (*Glycine max* (L.) Merrill) [13–15]. These few studies lead to the following question: How preserved is this spatial pattern across Families and within Families?

Additionally, it is worth highlighting that most nutrient studies in leguminous seeds relate to their total content. However, studies regarding nutrient distribution on the seed are scarce, yet crucial for a better understanding of the mechanisms underlying the seed physiology [16].

In this context, the microprobe X-ray fluorescence spectroscopy (µ-XRF) is an analytical technique suitable to study the nutrient composition and distribution of biological sample materials, including seeds [12,17–20] under *vivo* conditions or with minimal sample preparation [20–23]. Previous studies uncovered the elemental distribution in developing soybean seeds, using u-XRF to analyse the remobilization of elemental nutrients during germination and early seedling development [16], as well as to investigate the composition of biofortified and developing rice (*Oryza sativa* (L.) seeds [17,18]. However, the distribution of nutrients at the tissue level remains unexplored across seeds from the Fabaceae family.

Herein, we explored a quantitative XRF approach to assess the similarity and variances of macro (K, P, S, and Ca) and micronutrients (Fe, Zn, and Mn) along with the three important legume species seeds, *i*.*e*., cowpea (*Vigna unguiculata* L.), kidney bean (*Phaseolus vulgaris* L.), and soybean (*Glycine max* L.).

## 2. METHODS

### 2.1. Sample preparation

Seeds of cowpea (*Vigna unguiculata* (L.) Walp. variety BRS-Xiquexique, Brazil), kidney bean (*Phaseolus vulgaris* (L.) variety IPR Campos Gerais, Brazil), and soybean (*Glycine max* (L.) Merrill variety M6410 IPRO, Lagoa Bonita, Brazil) were manually cross-sectioned at the median region using a stainless-steel razor blade, preserving the embryo region. Afterwards, the sampled seeds were externally and individually fixed on 6 µm thick polypropylene film (VHG, FPPP25-R3 USA) mounted in an X-ray sample cup (Chemplex no. 1530, USA) for the XRF analyses. The experiments were carried out using three independent biological replicates, as detailed in Fig. S1.

### 2.2. Space-resolved X-ray fluorescence analysis

The sampled seed cross-sections were measured by μ-XRF (Orbis PC EDAX, USA). The elemental distribution of the macronutrients sulphur (S), phosphorus (P), calcium (Ca), and potassium (K), were investigated using a 64 × 50 pixel (n=3200 points) matrix, whereas the micronutrients zinc (Zn), manganese (Mn), and iron (Fe), distribution were determined throughout three 64-point line scanning across the embryo and cotyledon regions as shown in Figure S1. Each point was investigated using a 30 µm wide X-ray beam upcoming from a 50W Rh anode operating at 20 kV and 200 µA during 2 s for the macronutrient measurements, and 40 kV and 350 with a 25 µm thick titanium (Ti) primary filter during 10 s for the micronutrients. The analyses were performed under a vacuum atmosphere (< 50 Torr), and the X-ray spectra were recorded by a silicon drift detector (SDD), with a dead time smaller than 10%.

### 2.3. Quantification strategy

The concentration of macronutrients, i.e., P, S, Ca, and K in the sampled seeds was determined using cellulose-based standards prepared with standard-grade analytical reagents (KNO_3_, Vetec, Brazil; NaH_2_PO_4_.H_2_O, Baker, USA; Ca(NO_3_)_2_.4H_2_O, Synth, Brazil; MgSO_4_.7H_2_O, Êxodo Científica, Brazil. The resulting 0.5 g cellulose (cellulose binder, particle size < 20 μm, Spex, USA) pellets were then analysed through 16-point µ-XRF linescans at the same analytical conditions employed for assessing the seeds. The calibration curves are presented in supplementary Fig. S1. The trueness of the method was evaluated using certified reference materials (NIST1573a, NIST1515, NIST1547, and IAEA-V-10, respectively), and recovery values for all elements were within a 104-120% range.

Due to the inhomogeneous thickness variation across seed tissues (roughly from tenths to 3 millimetres), the pellet cellulose-based external calibration was not a suitable strategy for quantification of micronutrients *i*.*e*., Mn, Fe, and Zn. Particularly, the seed thickness range is under intermediate thickness XRF condition for the latter nutrients, in which distinct thickness directly affects the nutrient X-ray fluorescence intensity, and consequently their content determination. Therefore, their recorded XRF elemental intensities were herein explored.

### 2.4. Data processing

Only the X-ray characteristic intensities above the instrumental limit of detection (ILOD) were considered valid. The ILOD was calculated using Eq. 1.

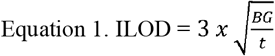

Where: BG (cps) is the average of 10 random measurements of the elemental background counting rate and t (s) is the acquisition time.

The spatial correlation between the distribution of macro and micronutrients was determined through Spearman’s correlation coefficient test at a 95% confidence interval, followed by a K-means clustering analysis. Besides, either the elemental concentrations or intensities recorded across each sampled seed tissue, i.e., seed coat, cotyledon, embryonal axis, and plumule, and the Krustall-Wallis analysis of variance followed by Dunn’s test at a fixed 0.05 significance level was carried out. All analyses were conducted using QtiPlot (version 5.9.8, Netherlands), RStudio (version 1.4.1106 “Tiger Daylily”, USA; packages ‘*stats*’ and ‘*ComplexHeatmap*’), and Prism (version 9.2.0, USA) software’s.

## 3. RESULTS AND DISCUSSION

### 3.1. Quantitative assessment of macronutrient distribution in Fabaceae seeds

To safeguard the generation renewal of the species, seeds do exhibit unique and highly conserved structural features for enabling their proper germination and early seedling development. In eudicots species, the seeds are usually coated by layers of parenchyma and sclerenchyma cells, in which a hilum, *i*.*e*., a scar left from the seed-funiculus attachment, stands out [24]. Internally, the seed is filled up mostly by cotyledons that encompass 90% of the area, with the rest of the embryo composed of a shoot apical meristem with an already emerging leaf primordium, *i*.*e*., plumule and radicular meristem, *i*.*e*., radicle, divided by the hypocotyl axis.

Figure 1 presents the quantitative microprobe XRF maps of P, S, K, and Ca in cowpea, soybean, and kidney bean seed cross-sections. The elemental images reveal that the spatial distribution of macronutrients is heterogeneous across seed tissues but does present conserved patterns among the three Fabaceae species. Similarly, Figures S2 and S2 show that the same trends were recorded in independent biological replicates.

**Figure 1.**
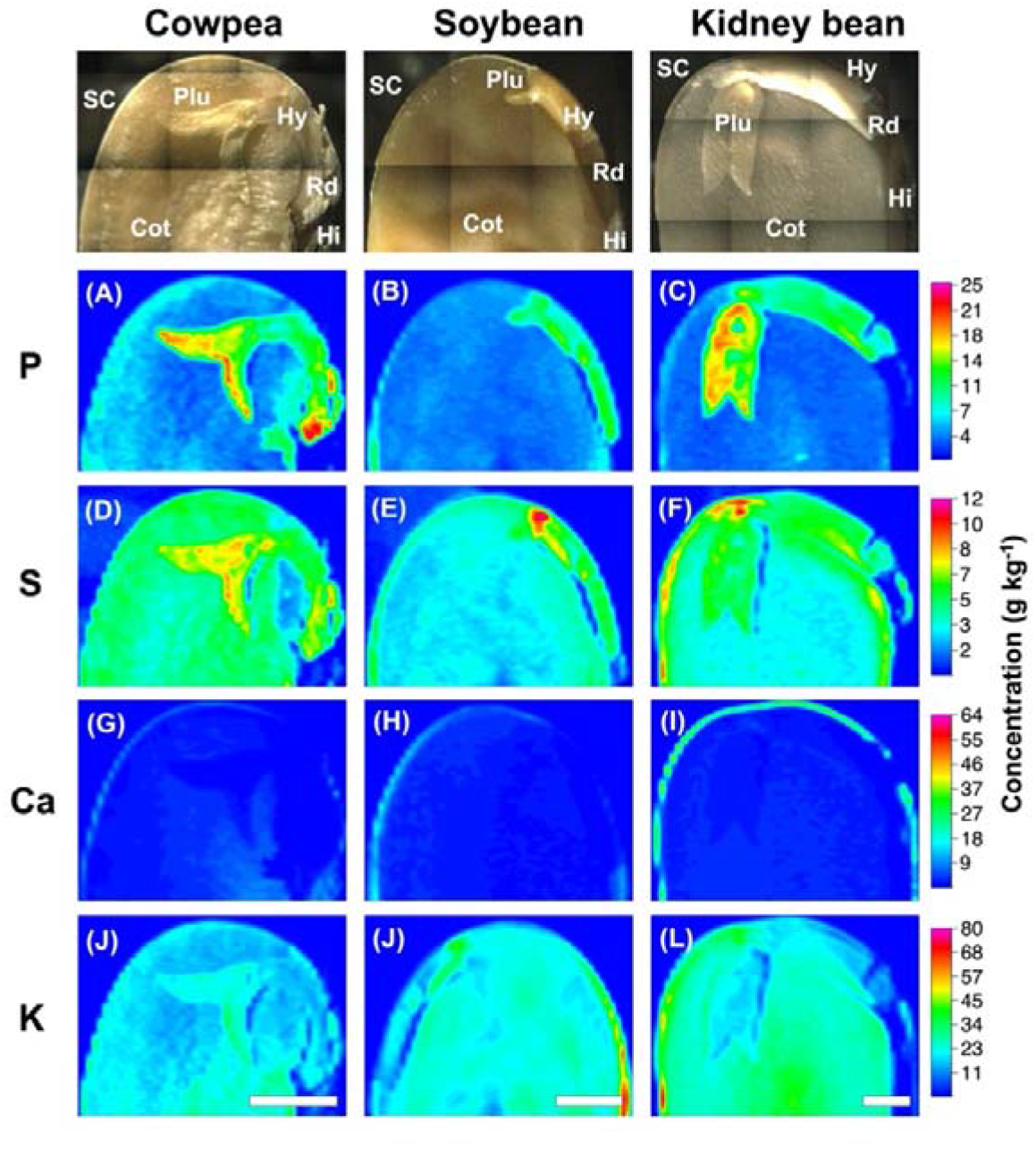
Quantitative XRF maps revealing the distribution of P (A-C), S (D-F), Ca (G-I), and K (J-L) in cowpea, soybean, and kidney bean seed cross-sections. The concentrations were determined through external cellulose-based calibration curves. SC: seed coat; Cot: cotyledon; Plu: plumule; Hy: hypocotyl; Rd: radicle; Hi: hilum. Scale: 2 mm.

Regardless of the species, one should notice that both P and S were mainly found in the embryos of the seeds, encompassing the embryonal axis, *i*.*e*., hypocotyl, radicle and plumule (Fig. 1 A-F). Conversely, Ca was located trapped in the seed coat and hilum (Fig. 1 G-I), the outermost seed tissues which play important roles either in the protection of seed structural integrity and water uptake during the very first stages of the germination process [25]. Interestingly, K was distributed throughout the whole seeds, with some hotspots surrounding either embryo or seed coat tissues (Fig. 1 J-L).

Herein, the spatial distributions for both Ca and K recorded in cowpea, soybean, and kidney bean seeds were akin to those found in the cross-sections of *Crotalaria ochroleuca* seeds [11], another specie from the Fabaceae family. Furthermore, Ca was also found mostly trapped in the tegument of rice (*Oryza sativa*) seeds [26], suggesting that seeds’ nutrient distribution patterns are highly conserved. However, it is important to highlight that these later studies explored a qualitative-based approach, thereby, the nutrient concentrations were not evaluated.

In this scenario, Table 1 presents the concentration of macro and micronutrients commonly found in the three species herein explored. Besides, Figure 2 presents the distribution of macronutrients found either in the seed coat, cotyledon, embryonal axis, and plumule seed tissues, and shows that P concentration in the seeds varied from 2-25 g kg^-1^ (Fig. 2 A-D). Based on these numbers, one can conclude that most of the P remains in the embryo axis and plumule tissues, whereas a minor fraction was observed in the cotyledon. Quantification of wheat seeds found P at 72 g kg^-1^ in the aleurone layer and 40 g kg^-1^ in parts of the embryo [34]. The high phosphorus concentration in the embryo axis and plumule can be associated with different tissues located in these embryo regions, considering the embryo axis is mostly composed of the non-differentiated primary meristems such as procambium, ground meristem, and protodermis [35]. In addition, the intense metabolic activity in meristematic cells is related to the high amount of P in the embryo’s axis and plumule cells.

**Table 1.** Mean contents of P, S, K, Ca, Mn, Fe, and Zn in soybean seeds (*Glycine max*), kidney bean (*Phaseolus vulgaris*), and cowpea (*Vigna unguiculata*).

|  | Element | Kidney bean | Cowpea | Soybean | Reference |
| --- | --- | --- | --- | --- | --- |
| Macronutrients<br>(g kg<sup>-1</sup>) | P | 5.5 | 4.7 | 10 | [27–29] |
|  | S | 11.6 | 2.4 | 5.4 | [27–29] |
|  | K | 38.5 | 1.1 | 20 | [27,29] |
|  | Ca | 26.9 | 0.7 | 30 | [27,29,30] |
| Micronutrients<br>(mg kg<sup>-1</sup>) | Zn | 27.61 | 45 | 42 | [30,31] |
|  | Fe | 40.6 | 50 | 71.5 | [28,31,32] |
|  | Mn | 16.5 | 15 | 30 | [30,31,33] |

**Figure 2.**
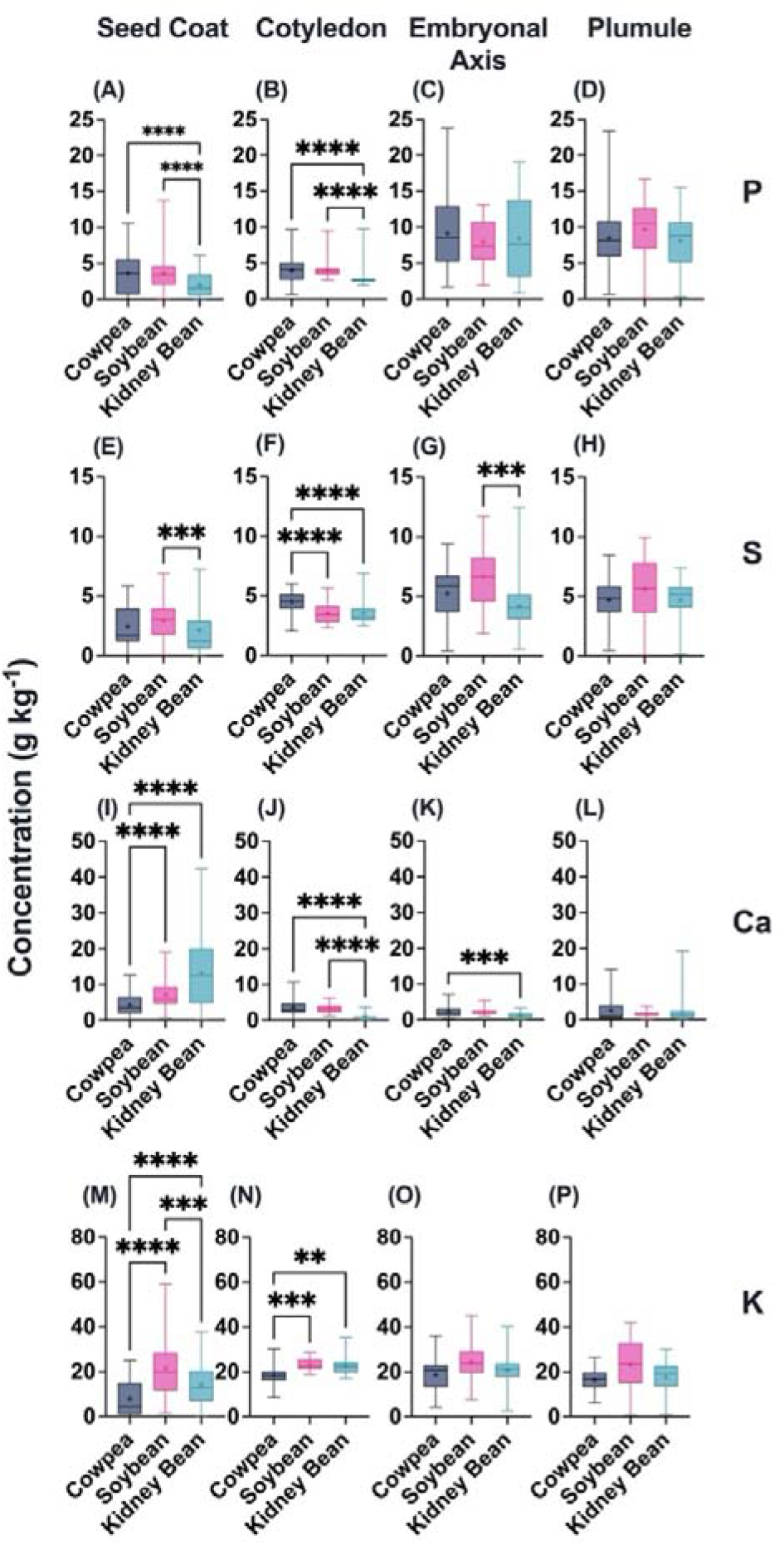
Boxplots of phosphorus (A-D), sulphur (E-H), calcium (I-L), and potassium (M-P) concentrations recorded at the seed coat, cotyledon, embryonal axis, and plumule tissues of cowpea, soybean, and kidney bean seed cross-sections. Data derived from three independent biological replications and were subjected to Krustall-Wallis’s analysis of variance followed by Dunn’s test at a fixed 0.05 significance level. (+) mean values; (*) p < 0.05), (**) p < 0.01, (***, ****) p < 0.001.

Furthermore, S concentrations ranged from 1 up to *ca*. 12 g kg^-1^ (Fig. 2 E-H) and were found mainly at the plumule for the three species. The highest S contents were found in the embryo of cowpea seeds and the peripheral zone of the cotyledon in the kidney bean. The S accumulation in the cowpea plumule can be explained by its well-developed leaf primordia. The seed species can be classified considering the predominant reserve in each species, therefore, soybean is classified as an aleuro-oilseed, storing mostly lipids and proteins, whereas beans are classified as aleurone-starchy, storing mostly protein and starch [36]. Sulphur is an element related to the essential amino acids such as cysteine and methionine that are presented as globular proteins associated with the protein storage vesicles in the cotyledonary parenchyma cells [37]. In the embryo axis, S may be part of amino acid molecules or coenzymes of lipid metabolite routes and the secondary <u>m</u>etabolism [38].

On the other hand, Ca was herein found mostly trapped in the tegument tissues, *i*.*e*., seed coat and hilum (Fig. 2 I-L), in which concentrations ranged from 1 up to 50 mg kg^-1^. The high Ca concentration in the tegument might be associated with the presence of a pectin-rich cell wall, mainly composed of Ca pectate, a Ca binding to non-methyl esterified homogalacturonan that affects the stiffness of the wall [39]. As the outermost tissues, both seed coat and hilum play crucial roles in protecting the seeds and modulating their interactions with the surrounding environment, *e*.*g*., water absorption that triggers the germination process [25]. Moreover, Ca is also located at plant cell vacuoles as prismatic crystals of Ca oxalate [40], but at a lower concentration. Besides, the µ-XRF analysis demonstrated that the Ca was also accumulated in the three studied species in the hilar region, similarly to the results recorded for soybean a in a previous XRF-based study [16].

Until now, the role of Ca at the hilum is not completely understood. Considering the water enters majorly through the hilum during the imbibition process, it is not clear whether the Ca deposited at the hilum can be remobilized to the embryo and then induce embryo germination.

Conversely, K concentration ranged from 1 to 52 g kg^-1^ in cowpea and varied from 1 to ca. 90 g kg^-1^ in kidney bean and soybean seeds (Fig. 2 M-P). Potassium plays an important role in the homeostasis of osmotic potential and ergastic substance biosynthesis, such as carbohydrates, as well as ensuring protein’s stability [41]. Besides, it was also verified K are easily remobilised from the cotyledonary parenchyma cells to the embryo axis as the primary roots emerge during hypocotyl elongation of the germination process of soybean seeds [16].

Furthermore, one should keep in mind that nutrient content in the seed might be related to its vigour. For example, Cotrim et al. found a positive correlation between the vigour and P, S, K, and Ca content along the embryonic axis of the *Crotalaria ochroleuca* seeds [11]. Then, the nutrient monitoring in seeds might also be explored as a strategy for evaluating their vigour.

### 3.2. Qualitative assessment of micronutrient distribution in Fabaceae seeds

For evaluating the spatial distribution of micronutrients in seeds, the linescan analysis approach showed to be more adequate than the maps, since the lower number of probed points enables exploring longer measurement time, thereby increasing the signal-to-noise ratio and the detection of trace elements, such as Mn, Fe, and Zn in vegetal samples [13,42–44]. In this regard, a triple series of 64-point line scans were performed for each triplicate of Fabaceae seeds. Supplementary Figure S4 details the linescans performed on seeds, while Figures S5-S7 show the XRF signals detected for Mn, Fe, and Zn in each one of the probed points.

Figure 3 presents the distribution of Mn, Fe, and Zn recorded either in the seed coat, cotyledon, embryonal axis, and plumule seed tissues. It reveals that Mn presented the lower XRF intensities compared to Fe and Zn, with the Mn values found in the cotyledon and embryonal axis tissues of soybean seeds (Fig. 3 A-D). Moreover, the Fe intensities were the highest among the micronutrients, also found mostly concentrated within the embryo axis. Aside from the embryo axis, a similar Fe concentration content was found in the cotyledon of all seed species (Fig. 3 E-H). Finally, the highest Zn XRF intensities were found in each bean-type embryo axis. Note that no statistically significant difference was found between the concentration of Zn in the seed coat and embryo among the beans (Fig. 3 I-L). On the other hand, Zn concentrations in the cotyledon and plumule were similar among the three species.

**Figure 3.**
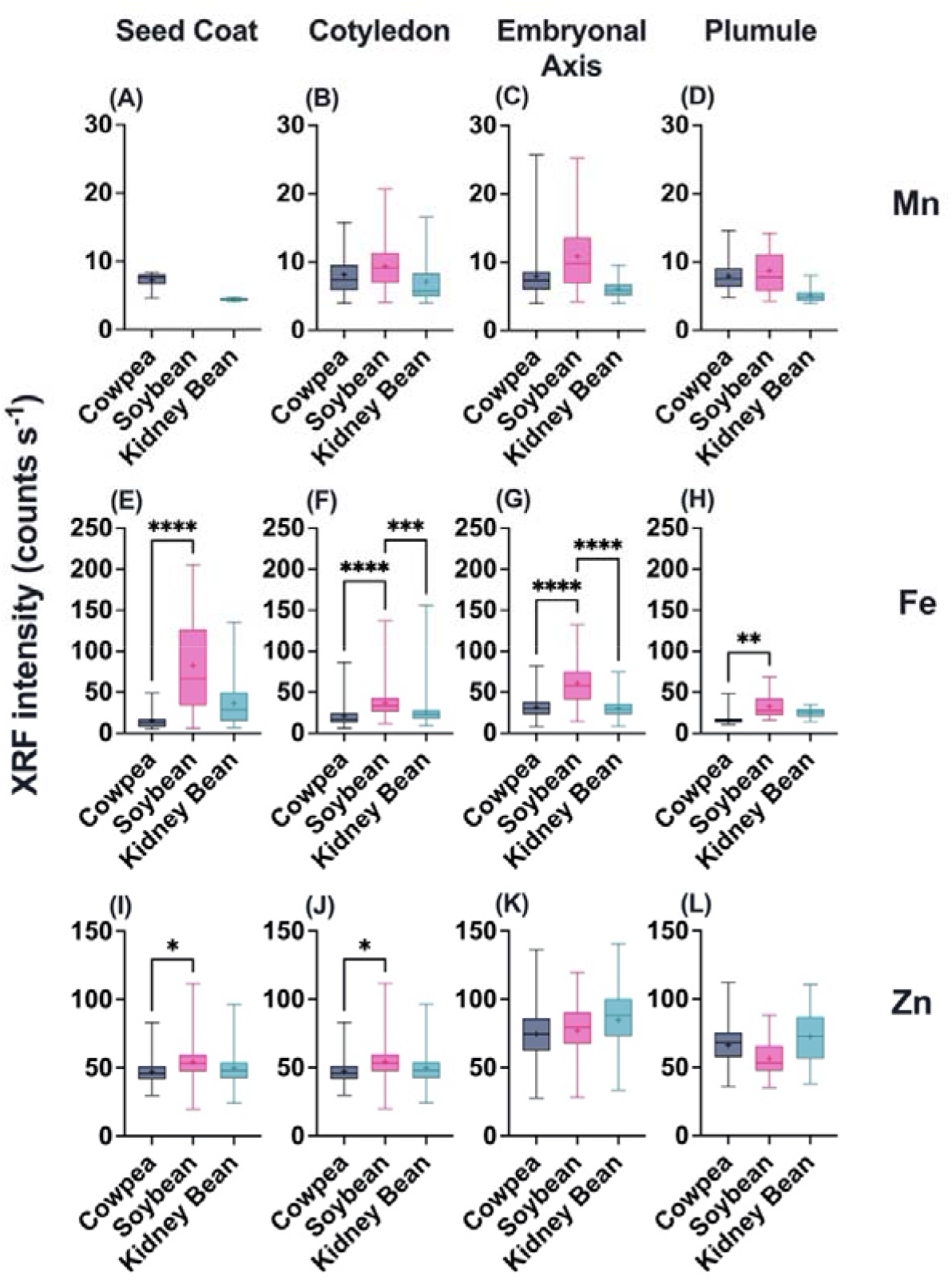
Boxplots of manganese (A-D), iron (E-H), and zinc (I-L) XRF elemental intensities recorded at the seed coat, cotyledon, embryonal axis, and plumule tissues of cowpea, soybean, and kidney bean seed cross-sections. Data derived from three independent biological replications and were subjected to Krustall-Wallis’s analysis of variance followed by Dunn’s test at a fixed 0.05 significance level. (+) mean values; (*) p < 0.05), (**) p < 0.01, (***, ****) p <0.001.

Synchrotron-based XRF maps of *Arabidopsis thaliana* showed a slightly higher Zn concentration of Zn in the cotyledon tissue compared to the radicle one [12]. In addition, as a cofactor for several enzymes related to gene expression, *e*.*g*., the Zn-finger transcriptional factor, the high amount of Zn found at the embryo axis and plumule might likely be associated with its role during seed germination and early development.

Furthermore, space-resolved microchemical maps of germinating rice grains cross-sections showed that Mn can be accumulated in the seed coat and scutellum, and then translocated to the leaf primordium [45]. Manganese participates in the metalloenzyme cluster of the oxygen-evolving complex in photosystem II, it acts as a cofactor for several enzymes related to the secondary metabolism and as enzymes that are associated with the detoxification process. Oxalate oxidase, for instance, is an Mn-dependent [46] highly expressed during the embryo germination of wheat, and likely plays an important role in the protection against oxidative stresses during seed maturation and germination [47]. In addition, Mn is also related to the formation of lignin that forms the cell wall [48].

Due to the low energy X-ray fluorescence absorption by the X-ray detector window and the low fluorescence yield, some lower atomic number macronutrients, *e*.*g*. B, Mg and N, were not detected. Additionally, the detection of some micronutrients, such as Cu and Mo, was not possible under the current instrumental conditions due to their lower concentration.

### 3.3. Tissue-resolved correlation of nutrients distribution in Fabaceae seeds

Aiming at understanding the spatial distribution patterns displayed by the probed elements across the Fabaceae seeds, Figure 4 presents the heatmaps of Spearman’s correlation coefficients determined across cowpea, soybean, and kidney bean seeds XRF recorded values.

**Figure 4.**
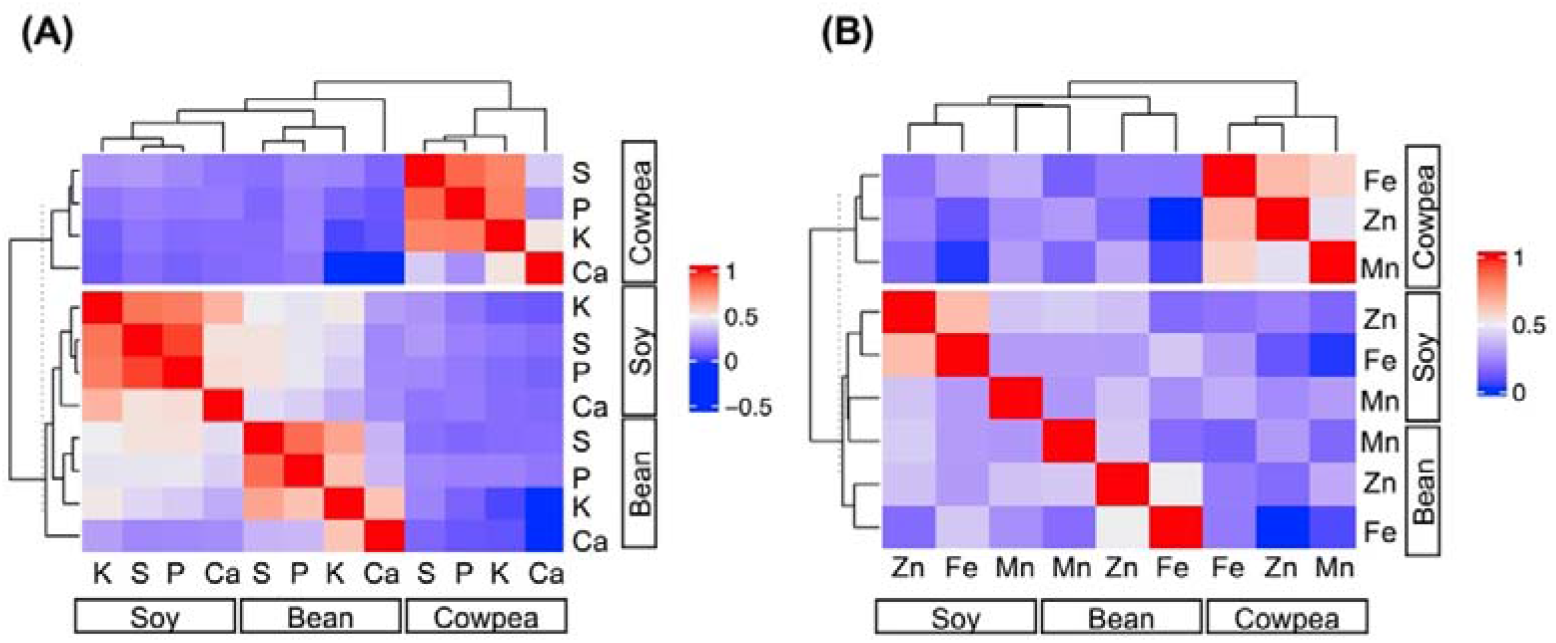
Heatmaps of Spearman’s correlations coefficient for XRF-probed macro (A) and micronutrients (B) spatial distribution across cowpea, soybean, and kidney bean cross-sections. The K-means clustering displays the similarity of the correlation values found for each seed species assessed.

For the macronutrients (Fig. 4 A), it reveals that the highest correlations were observed between P and S, S and K. Conversely, Ca presented the lowest correlations in the three Fabaceae, showing lower correlations with K in cowpea, both K and P in the kidney bean ones, and with S in soybean seeds. These results follow a similar trend as those observed *in* developing soybean seeds [16]. For the micronutrients (Fig. 4 B), higher correlations values were usually recorded compared to the macronutrient ones. An interesting association between Fe and Mn, and Fe and Zn were observed in cowpea and soybean seeds, respectively. A similar association between Mn, Fe, and Zn was observed in the *Turkish pea* [49].

Furthermore, the K-means clustering also revealed that regardless of macro and micronutrients, the elemental distribution of kidney bean and soybean and soybean seeds were more related compared to the cowpea ones.

## 4. CONCLUSION

The present approach unveils the feasibility of the X-ray fluorescence spectroscopy to depict the elemental distribution of seeds, and unveils that the nutrients are not homogeneously distributed in Fabaceae seeds, but follow similar trends. Sulphur and P concentrated mostly in the seed’s embryo, whereas Ca was mostly in the seed coat and hilum, and K presented a more homogenous distribution throughout the seeds. Besides, Zn and Mn seem to share similar locations mainly throughout the embryo. These results enhance the understanding of the nutrient composition of seeds and might shed light on the understanding of their roles in seed physiology. This current approach might be useful for helping in the development of nutritional strategies that might enhance seed vigour and the performance of their upcoming plants in the field.

## ACKNOWLEDGEMENTS AND FUNDING

The X-ray spectrometry facilities used in this study were funded by São Paulo Research Foundation (FAPESP grant 2015/19121-8). G.S. Montanha is the recipient of a FAPESP doctoral scholarship (2020/07721-9). L.A. Rohr is the recipient of a FAPESP master’s degree scholarship (2019/17967-8). H.W.P. Carvalho is the recipient of a research productivity fellowship from the Brazilian National Council for Scientific and Technological Development (CNPq) (grant 306185/2020-2).

## AUTHORSHIP

**[Conceptualization]** H.W.P.C., J.P.R.M., and G.S.M.; **[Methodology]** H.W.P.C., S.L.Z.R., and G.S.M.; **[Resources]** H.W.P.C.; **[Data curation]** G.S.M, S.L.Z.R., and H.W.P.C.; **[Original Draft Preparation]** G.SM., S.L.Z.R., E.A., L.A.R., J.P.R.M., and H.W.P.C.; **[Review & Editing]** S.L.Z.R., G.SM., E.A., L.A.R., J.P.R.M., F.S.L., and H.W.P.C

## DATA AVAILABILITY STATEMENT

The data underlying this article will be shared on reasonable request to the corresponding author.

## ELECTRONIC SUPPLEMENTARY MATERIAL

**Figure S1.**
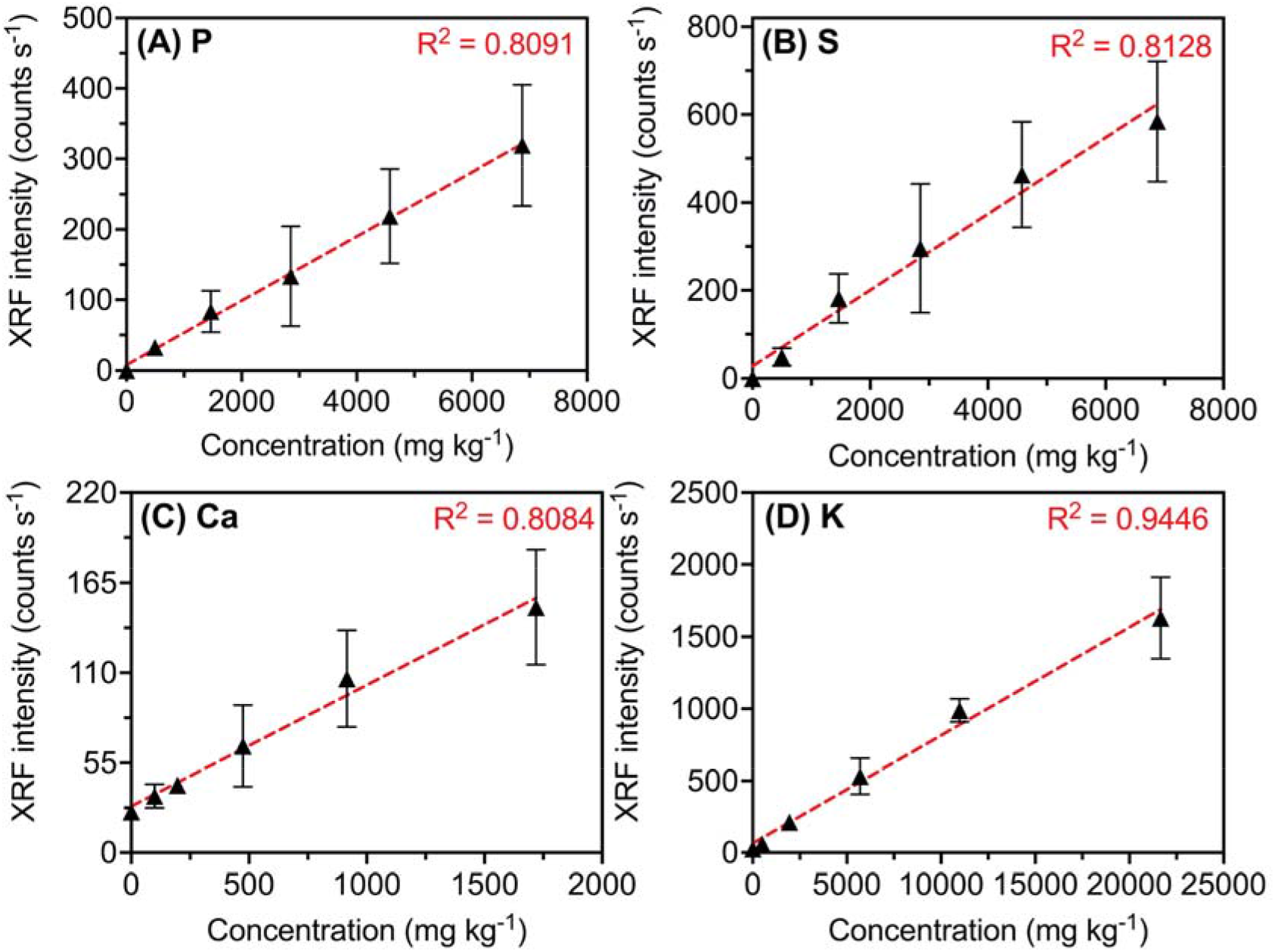
Calibration curves derived from cellulose-based standards for P, S, Ca, and K, respectively. Values derived from 16 independent measurements.

**Figure S2.**
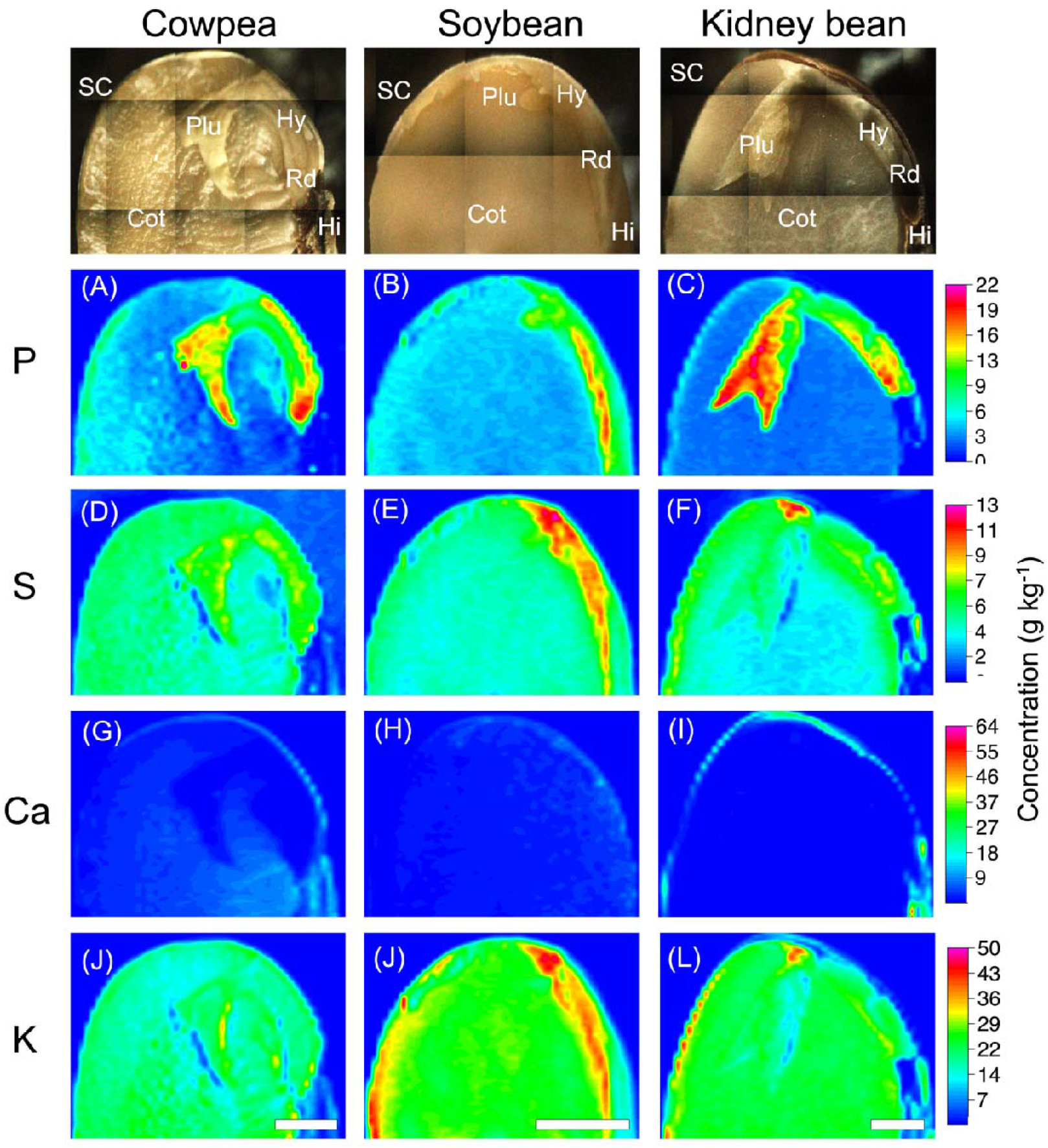
Quantitative XRF maps revealing the distribution of P (A-C), S (D-F), Ca (G-I), and K (J-L) in cowpea, soybean, and kidney bean seed cross-sections. The concentrations were determined through external cellulose-based calibration curves. SC: seed coat; Cot: cotyledon; Plu: plumule; Hy: hypocotyl; Rd: radicle; Hi: hilum. Scale: 2 mm. Data from an independent biological replicate.

**Figure S3.**
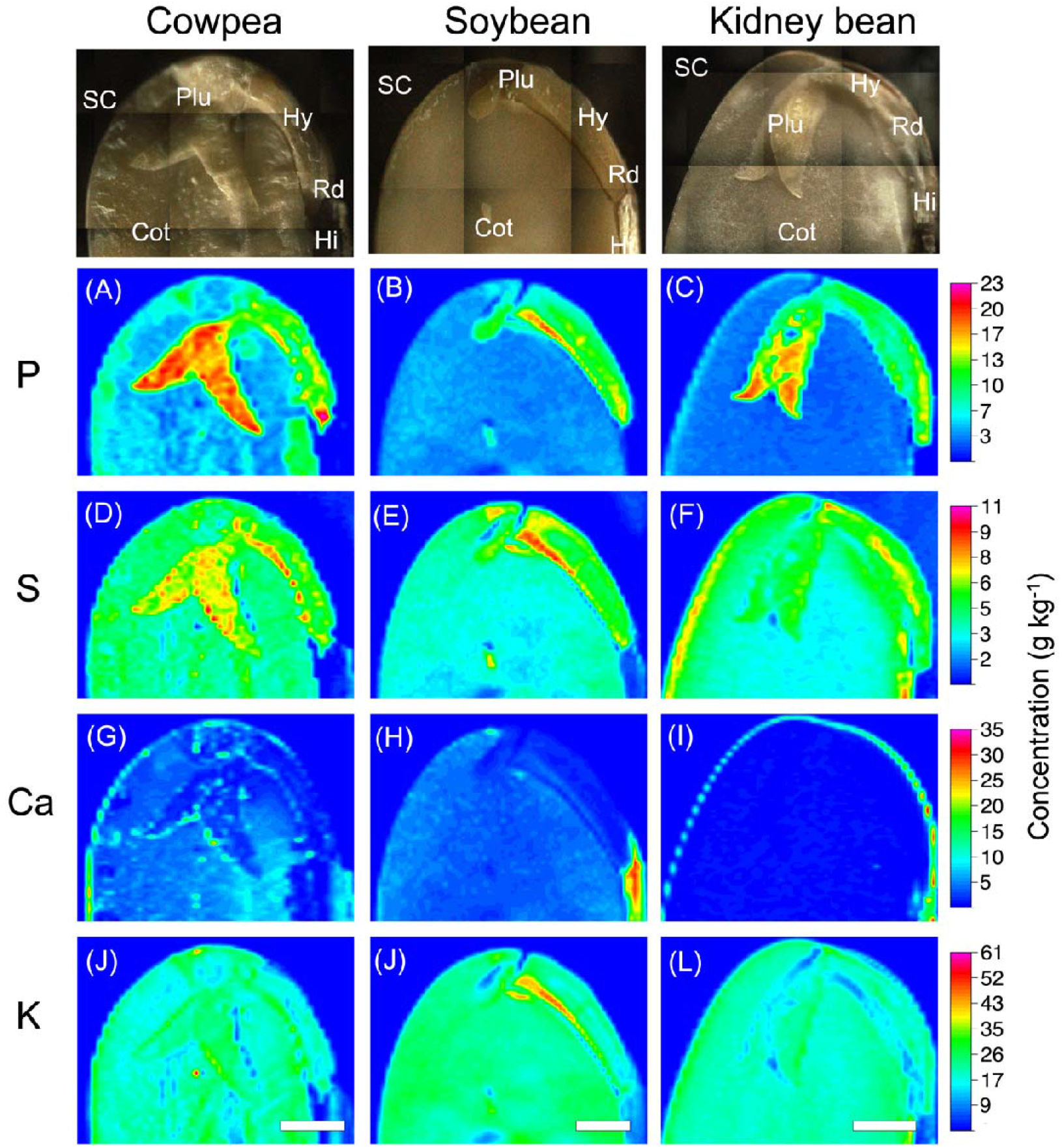
Quantitative XRF maps revealing the distribution of P (A-C), S (D-F), Ca (G-I), and K (J-L) in cowpea, soybean, and kidney bean seed cross-sections. The concentrations were determined through external cellulose-based calibration curves. SC: seed coat; Cot: cotyledon; Plu: plumule; Hy: hypocotyl; Rd: radicle; Hi: hilum. Scale: 2 mm. Data from an independent biological replicate.

**Figure S4.**
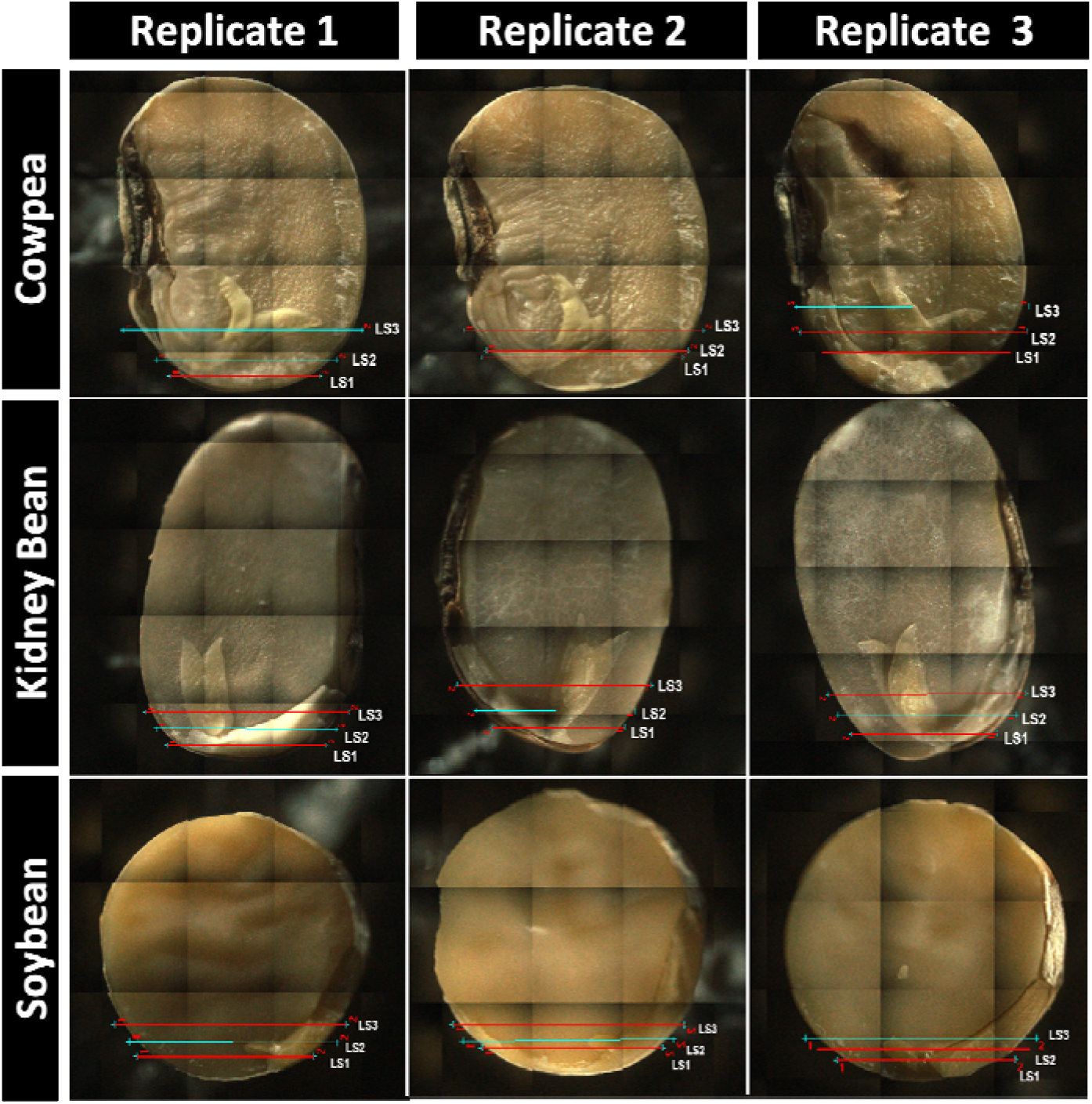
Photographs and representation of each line scanned through microprobe X-ray fluorescence spectroscopy to evaluate the distribution of micronutrients, e.g., Mn, Fe, and Zn found across the tissues of cowpea, soybean, and kidney bean seed cross-sections.

**Figure S5.**
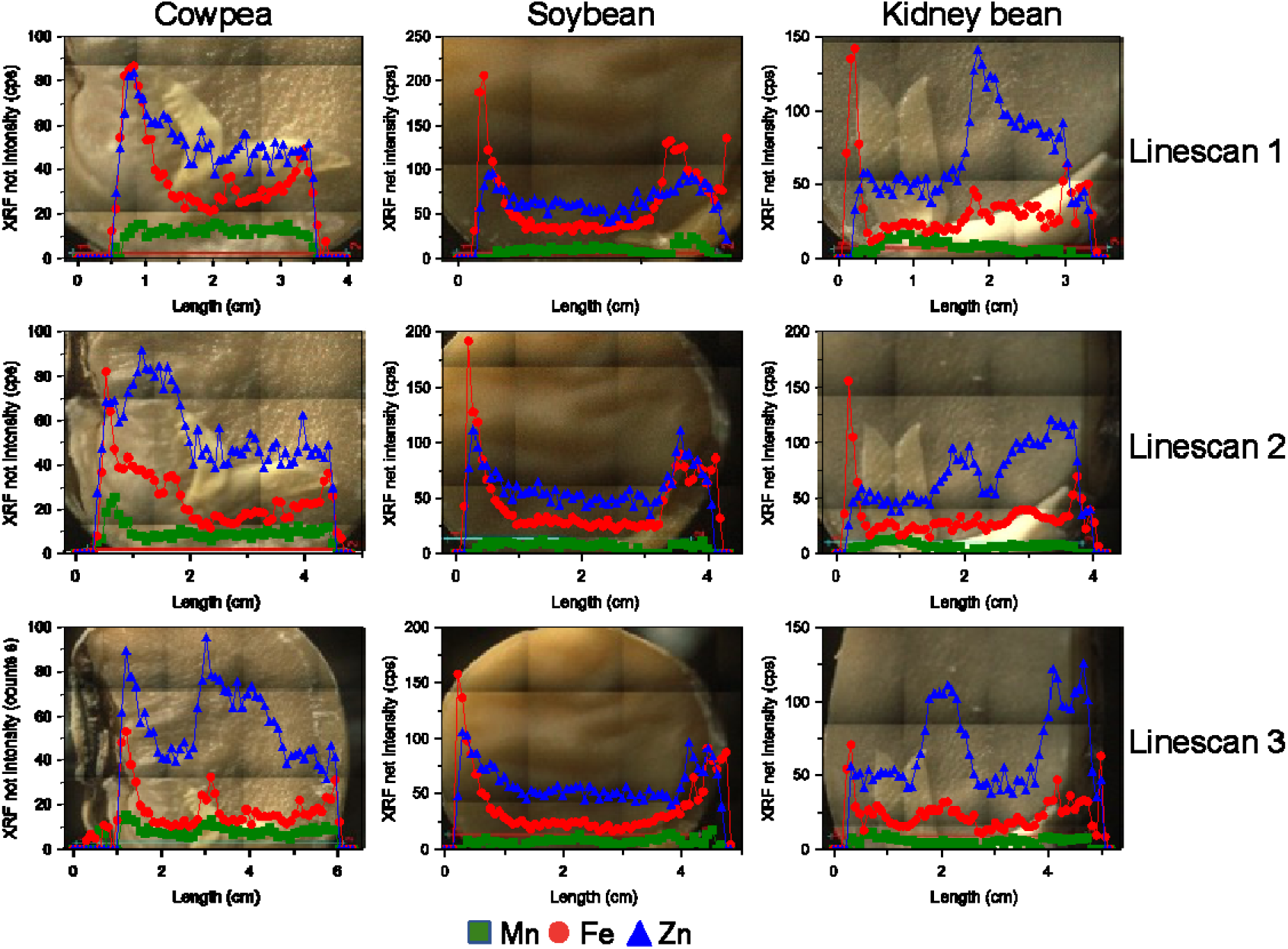
XRF probing Mn, Fe, and Zn spatial distribution along cowpea, soybean, and kidney bean seed cross-sections. Three independent linescans were recorded at each seed specie. Data from an independent biological replicate.

**Figure S6.**
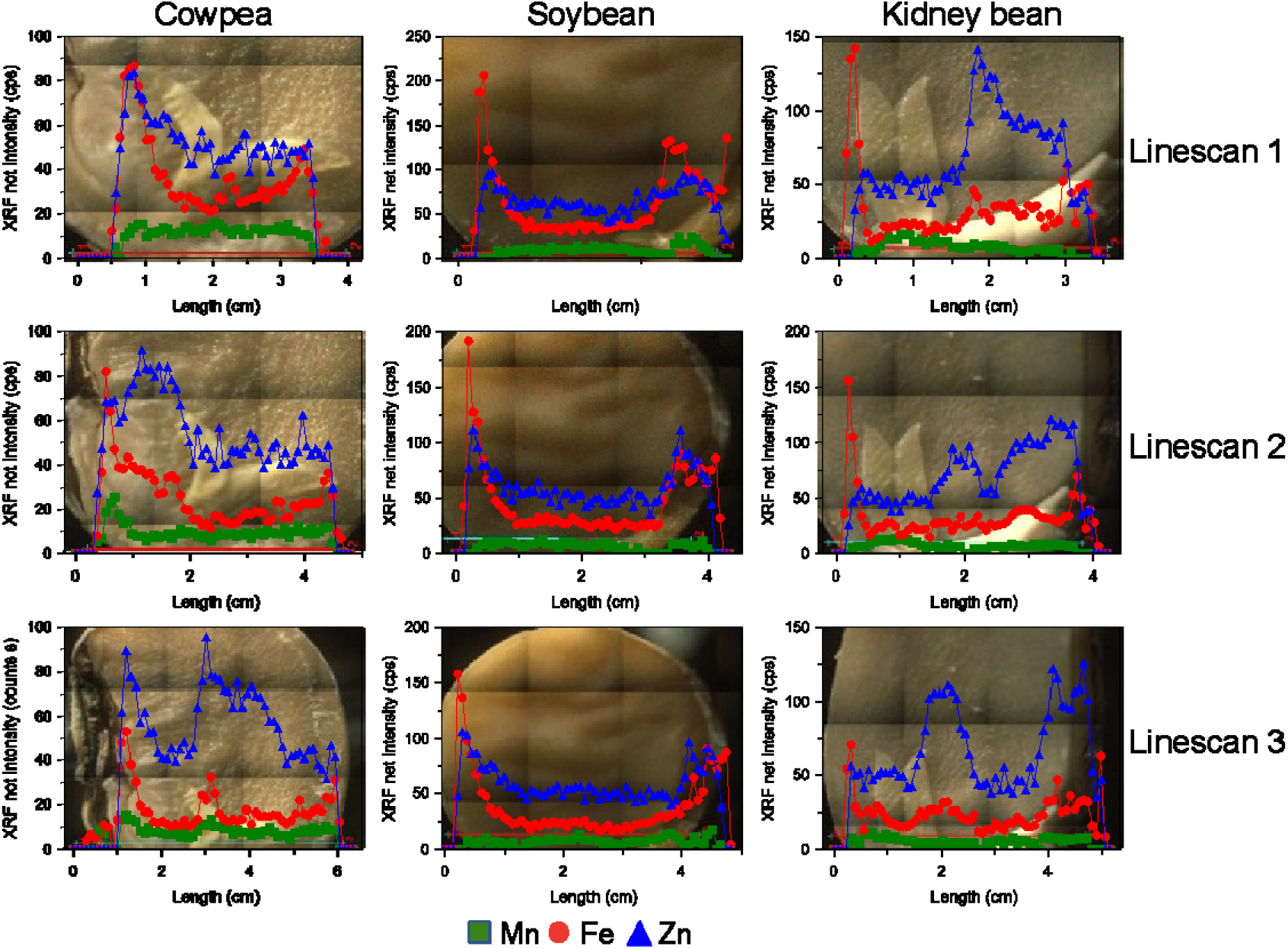
XRF probing Mn, Fe, and Zn spatial distribution along cowpea, soybean, and kidney bean seed cross-sections. Three independent linescans were recorded at each seed specie. Data from an independent biological replicate.

**Figure S7.**
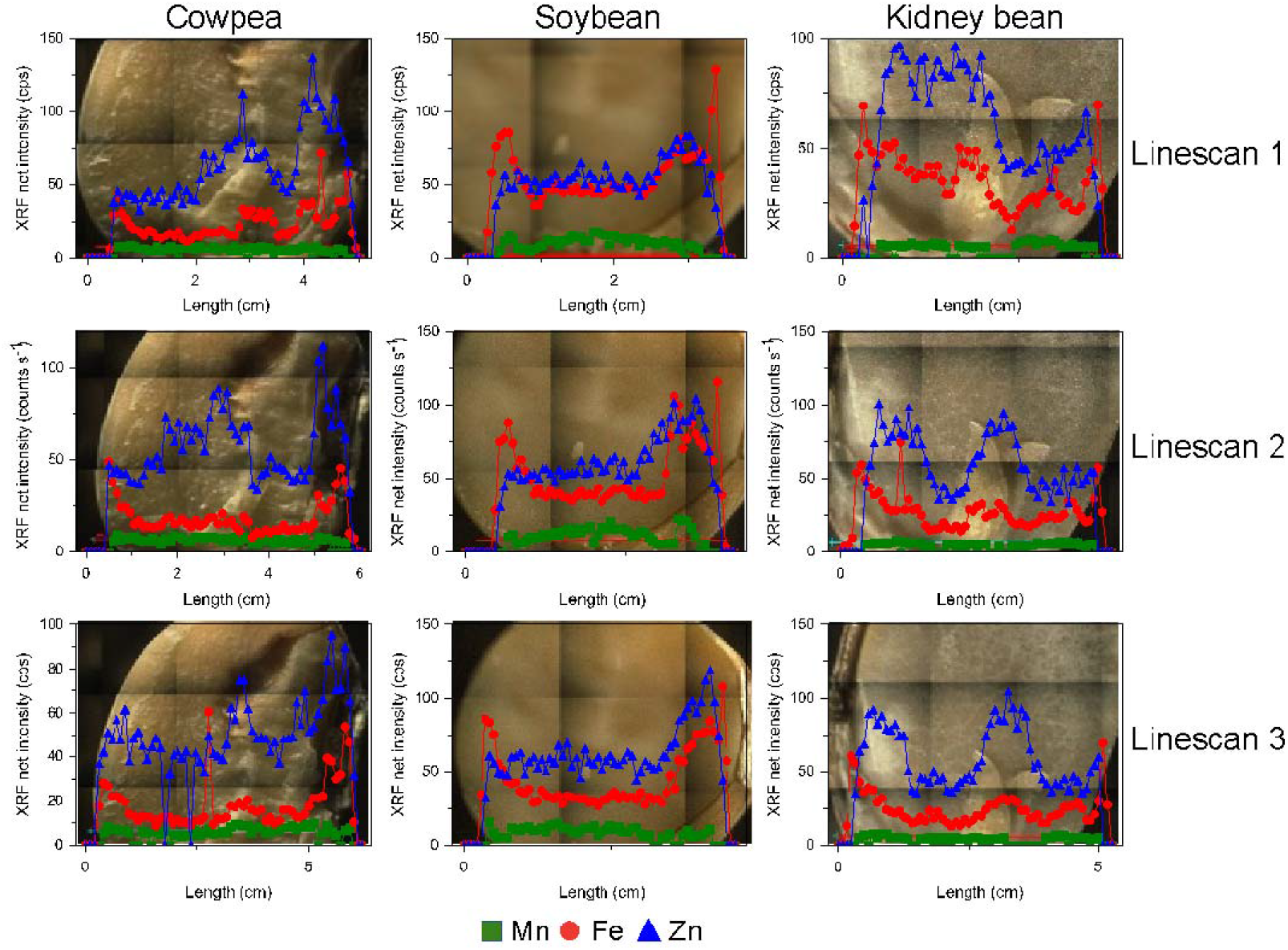
XRF probing Mn, Fe, and Zn spatial distribution along cowpea, soybean, and kidney bean seed cross-sections. Three independent linescans were recorded at each seed specie. Data from an independent biological replicate.

